# Summarizing Performance for Genome Scale Measurement of miRNA: Reference Samples and Metrics

**DOI:** 10.1101/210310

**Authors:** PS Pine, SP Lund, JR Parsons, LK Vang, AA Mahabal, L Cinquini, SC Kelly, H Kincaid, DJ Crichton, A Spira, G Liu, AC Gower, HI Pass, C Goparaju, SM Dubinett, K Krysan, SA Stass, D Kukuruga, K Van Keuren-Jensen, A Courtright-Lim, KL Thompson, BA Rosenzweig, L Sorbara, S Srivastava, ML Salit

## Abstract

**Background:** The potential utility of microRNA as biomarkers for early detection of cancer and other diseases is being investigated with genome-scale profiling of differentially expressed microRNA. Processes for measurement assurance are critical components of genome-scale measurements. Here, we evaluated the utility of a set of total RNA samples, designed with between-sample differences in the relative abundance of miRNAs, as process controls.

**Results:** Three pure total human RNA samples (brain, liver, and placenta) and two different mixtures of these components were evaluated as measurement assurance control samples on multiple measurement systems at multiple sites and over multiple rounds. *In silico* modeling of mixtures provided benchmark values for comparison with physical mixtures. Biomarker development laboratories using next-generation sequencing (NGS) or genome-scale hybridization assays participated in the study and returned data from the samples using their routine workflows. Multiplexed and single assay reverse-transcription PCR (RT-PCR) was used to confirm *in silico* predicted sample differences. Data visualizations and summary metrics for genome-scale miRNA profiling assessment were developed using this dataset, and a range of performance was observed. These metrics have been incorporated into an online data analysis pipeline and provide a convenient *dashboard* view of results from experiments following the described design. The website also serves as a repository for the accumulation of performance values providing new participants in the project an opportunity to learn what may be achievable with similar measurement processes.

**Conclusions:** The set of reference samples used in this study provides benchmark values suitable for assessing genome-scale miRNA profiling processes. Incorporation of these metrics into an online resource allows laboratories to periodically evaluate their performance and assess any changes introduced into their measurement process.

## Background

Studies to identify potential biomarkers typically involve comparing two conditions and identifying features that distinguish the two classes; for example, disease versus normal or treated versus control. Quality measurements made during this discovery phase are essential for the success of all subsequent phases of biomarker development. Reference samples, with known differences, can enable laboratories to assess and improve their ability to detect relevant biomarkers. Within the framework of the Early Detection Research Network (EDRN) of the National Cancer Institute [1], we are developing a measurement assurance paradigm for genome-scale measurement systems currently used for microRNA (miRNA) biomarker discovery.

Comparisons of the results for genome-scale measurements of two different biological samples or two different reference samples have been used to assess both microarray [2, 3] and RNA sequencing (RNAseq) measurements of messenger RNA (mRNA) [4]. Evaluations derived from this type of comparison are limited to metrics such as concordance of gene lists and correlations of rank order because, in both cases, the true difference between samples is not known. For both microarray and RNAseq, titration designs have also been used, which provide some information regarding signal trends [4, 5].

Composite reference samples with designed-in differences provide additional metrics for performance assessment and have been demonstrated to be useful with both microarrays [6 – 9] and RNAseq [10]. These same technologies have been applied more recently to profiling miRNA, a class of small non-protein-coding RNAs that regulate the expression of hundreds of target genes by translational repression, controlling biological functions involved in differentiation and development. Characterizing miRNA measurements on multiple platforms has been performed with biological samples [11] and titrations of biological samples [12]. Here we demonstrate the utility of one of these mixture designs (Figure 1) with corresponding metrics [7 – 10] and introduce new metrics useful for summarizing and evaluating genome-scale measurements developed in an interlaboratory study spanning multiple rounds of measurement. Multiple data visualizations and metrics have been combined into a single standardized view, or “dashboard”, for each participant and round (see Figure 2 for an example). These are available in Additional file 1. Representative panels are described in detail in the results.

**Figure 1.**
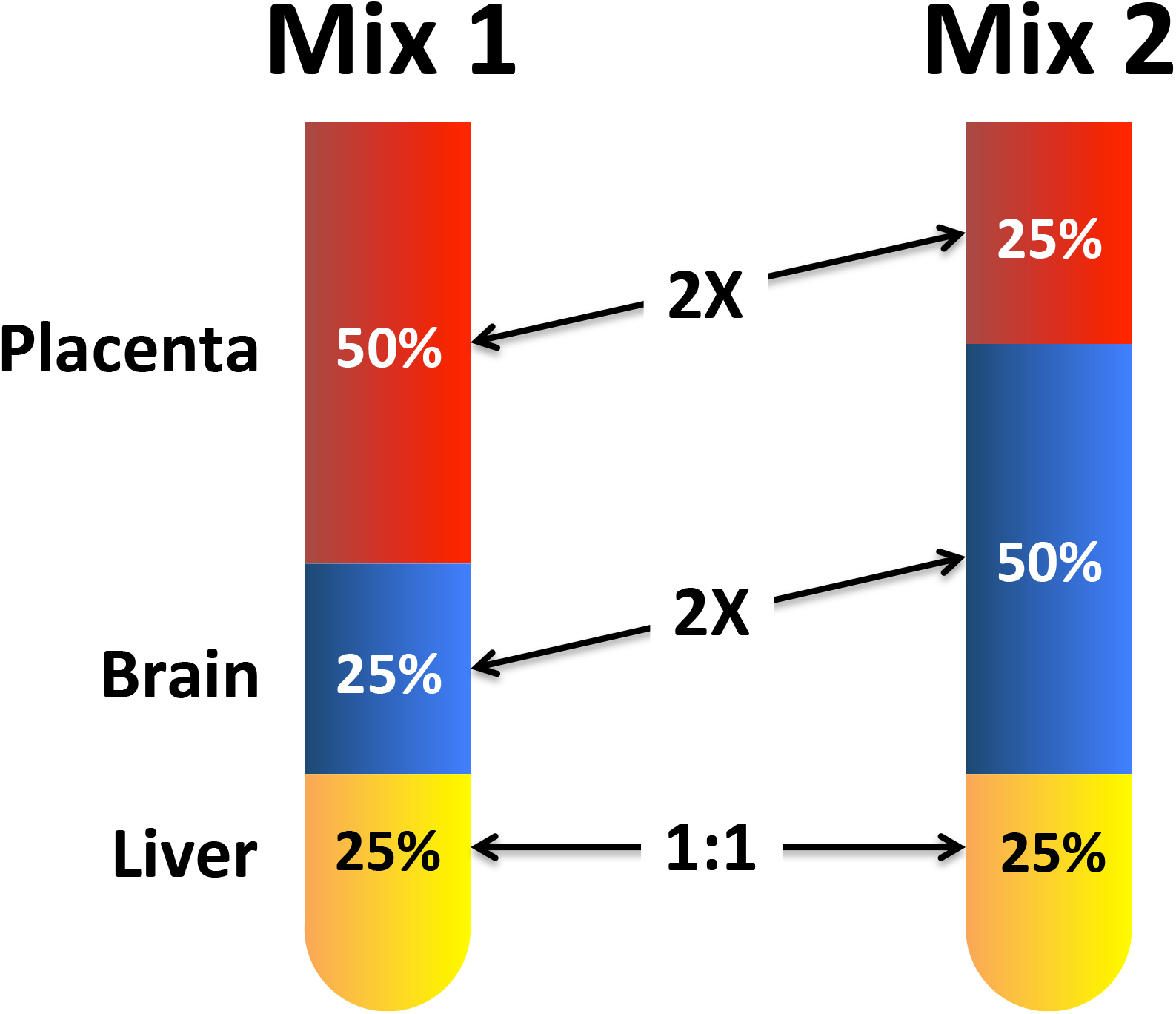
The relative input proportions from three total RNA components are shown for Mix1 and Mix2.

**Figure 2.**
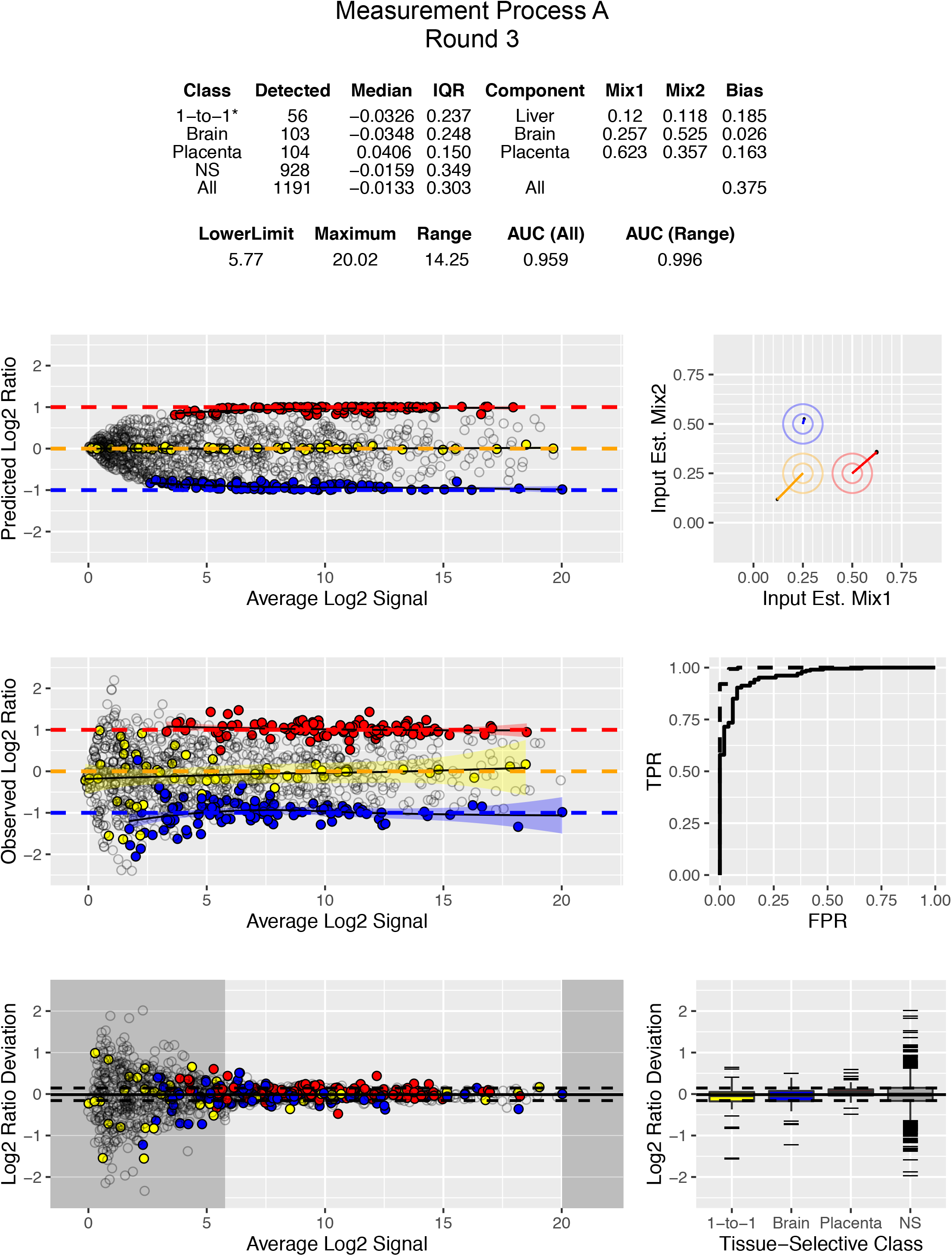
Dashboard view combining visualizations and metrics. Metrics table displays values derived from data visualizations. The legends for Figures 3 to 8 describe each panel.

## Results

### Selection of human tissue total RNA

Previous studies have demonstrated the feasibility of using a pair of reference mixtures with designed-in differences to provide performance benchmarks [6–10]. These rely on a combination of pure total RNA components with a large number of transcripts that are distributed across a broad dynamic range of relative abundance. Each component should contain a subset of miRNA that are either unique to that tissue or enriched relative to the other two components. These tissue-selective miRNAs will be used in metrics to assess assay performance. The tissue-selectivity can be quantified [7] and used to assess whether a sufficient number of differentially abundant miRNA are available in each subset to span the dynamic range of the measurement process. Laboratories performing biomarker discovery may prefer one of the components be similar in nature to the tissue being profiled in their own research.

To select components for reference mixtures described in this paper, a published study comparing the miRNA expression profiles of nine different tissues using both microarrays and RT-PCR was utilized [13]. The two sources of human RNA with the most tissue-selective content were placenta followed by brain. These two tissue RNAs were used as variable components in a reciprocal two-to-one design, with liver as the invariable (one-to-one) component (Figure 1) [14].

### In silico modeling

The miRNA signals in each mixture should be an additive and linear combination of the signals from pure tissue components [6 – 10]. Therefore, an expected signal and predicted ratio can be calculated based upon the fractional proportion of each tissue component in a mixture using the following equations:

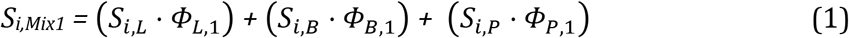

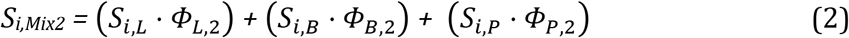

Equations 1 and 2 show the formulae for the pair of three component mixture designs, where *S* is the signal from a particular miRNA *i* and *ϕ* is the fraction of total RNA for each tissue (liver=*L*, brain=*B*, and placenta=*P*) in mixtures *1* and *2*. For example, using mixture proportions of 1:1:2 and 1:2:1 (L:B:P) provides corresponding *ϕ* of 0.25, 0.25, 0.5 and 0.25, 0.5, 0.25, respectively. With this design, the maximum possible ratio (*S*_*i,Mix1*_/*S*_*i,Mix2*_) for any miRNA in the final mixture comparison corresponds to a 2-fold difference (i.e., log2 difference between Mix1 and Mix2 of −1 or 1) which would be observed for brain-specific or placenta-specific miRNAs. For miRNAs that are not tissue-selective, some signals will be contributed from each component, resulting in ratios falling somewhere within that range. Estimating mixture signals from measured signals of unmixed tissues using *in silico* modeling with Equations 1 and 2 provides predicted values for comparison to observed results for Mix1 and Mix2.

### Ratio Estimates

The first two rounds of measurement (Rounds 1 and 2, not shown) were pilot studies used for tissue profiling alone, and did not include mixtures. A total of 7 sites participated in the three rounds (Rounds 3 to 5) that included the three pure RNA samples and the two mixtures of them. In each of these rounds, each site received three replicates for each of the 5 samples, with sample identities hidden. Participants profiled miRNA expression with the platforms used in their routine workflow (one site used two different platforms). Labs using genome-scale platforms reported all detectable miRNAs. Labs using RT-PCR performed assays for a subset of miRNAs of interest to confirm that the mixtures could produce observed ratios similar to predicted values.

Log2 transformed ratio estimates for miRNAs were calculated for each round and compared across multiple participants. For analysis of genome-scale data, detectable miRNAs (counts ≥ 1 in any one sample) were normalized to the median total count among the samples and then log2 transformed. However, raw count tables, as well as datasets preprocessed with other strategies, could also be used as input. For RT-PCR data, the quantitation cycle (Cq) values were negatively transformed to provide comparable log2 transformed data. Predicted log2 ratios were calculated from the pair-wise differences of the modeled mixtures derived from Equations 1 and 2 using the linear transformed means for each pure tissue component, *S*_*i*_. Observed log2 ratios were estimated from the pair-wise differences between means for each mixture, Mix1 and Mix2, described in Methods.

### Visualizations and analyses

For the sites using genome-scale technologies, the log2 ratios for all detectable miRNA can be visualized using a Bland-Altman plot [15] to evaluate the ratio data throughout the dynamic range, and any miRNA that are highly enriched in one tissue relative to the other tissues should approach the benchmark values of the mixture designs.

In Figure 3, the *predicted* ratios for all detected miRNA were calculated using Equations 1 and 2. The 10X selective miRNA (those miRNA that are at least 10 times more prevalent in one tissue relative to the others) are derived from comparisons of the profiles of the three pure RNA samples included in the sample set, and are color-coded according to tissue type as in Figure 1. Panels A to D represent multiple sites using different NGS platforms, with different sequencing depths. One site (panel A) was able to detect 1130 miRNA consistently across the 15 samples in the set, with 107 brain-selective and 105 placenta-selective miRNA with log2 ratios distinguishable from the “1-to-1” class, 59 miRNA with predicted log2 ratios of approximately zero (i.e., no difference between Mix1 and Mix2). The 1-to-1 class includes 23 liver-selective miRNA and 36 miRNA with approximately equal amounts of signal for brain and placenta. For much of the dynamic range the tissue-selective miRNA are predicted to approximate the designed-in log2 ratio limits of ± 1. In comparison, another site (panel D) using a different NGS platform producing fewer reads and detected 291 miRNA in total. Of those, only 18 brain-selective, 9 placenta-selective, and 16 miRNA predicted to be 1-to-1 were identified. Panel E shows the results from the site using a hybridization-based platform [16]. The large cluster of non-selective (NS) miRNA near the low end of the dynamic range is consistent with a background level of hybridization and this type of additive noise is known to contribute to ratio compression. One site (panel F, circles) used a multiplexed PCR platform targeting 32 different miRNA that were selected for their presence on common fixed content platforms [17]. Based on modeling of the data from the microarray study [13], two additional PCR sites were asked to measure a subset of miRNAs predicted to provide differences between mixtures: ~2 -fold up for miR-451a and miR-335; ~2 -fold down for miR-125b and miR-218; and no change for miR-375. One PCR site measured all five miRNAs in Round 3 (panel F, squares) and the other site measured two (panel F, triangles). Detection metrics are part of the summary table included in the dashboard view for each site and round (Additional file 1).

**Figure 3.**
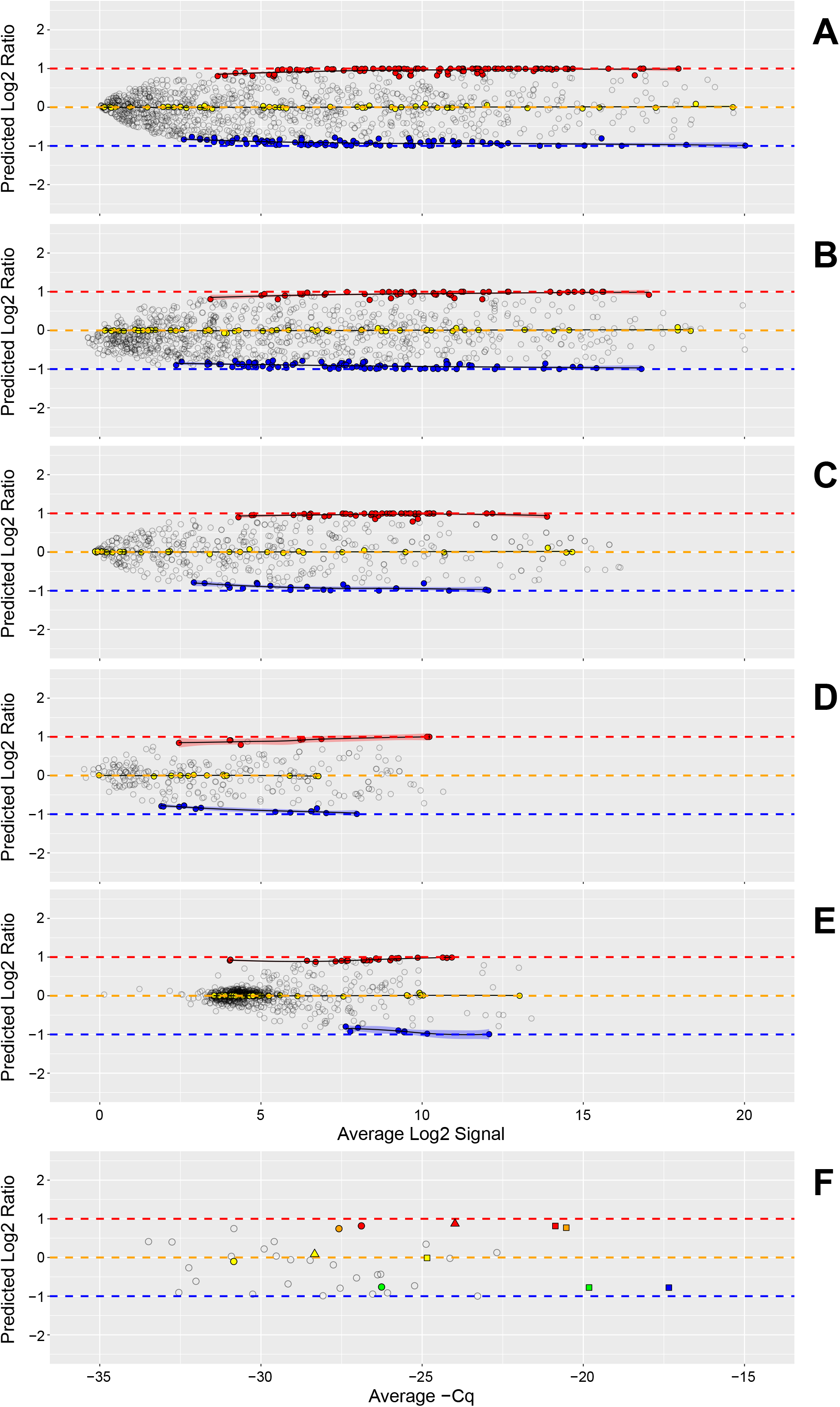
Predicted distribution of log2 ratios. Panels A to F correspond to measurement processes A to F described in Results. For each datapoint in panels A to F, the difference between the predicted Mix1 and Mix2 log2 signals (log2 ratios) is plotted against their average for each detected miRNA. Signal values for each mixture are predicted using Equations 1 and 2. Filled circles correspond to predicted values for miRNA that were at least 10-fold enriched in relative abundance compared with the other two tissues, or miRNA that were approximately equal in relative abundance between placenta and brain (1-to-1): red = 10X placenta, blue = 10X brain, and yellow = 1 -to-1 (10X liver and placenta = brain). Open circles correspond to detectable, but nonselective miRNA. Red, yellow, and blue transparent bands indicate the 95% confidence interval for the loess (locally weighted smoothing) function (black lines) for the placenta, 1 -to-1, and brain subsets, respectively. Panel F includes data from three different PCR labs: one site using multiplexed PCR (circles) and two sites using individual PCR assays (squares and triangles). Five miRNA of interest are highlighted: miR-451a (red), miR-335 (orange), miR-375 (yellow), miR-218 (green), miR-125b (blue). *The total number of detectable miRNA and their tissue-selective classification are included in the summary table of the dashboard.*

Figure 4 shows the experimentally *observed* ratios for miRNAs in Mix1 and Mix2 (see Figure 1 for composition) and corresponds to the same labs displayed in Figure 3, panels A to F. There is more dispersion in the observed log2 ratio data when compared to the predicted values, which also becomes more apparent at the lower end of the dynamic range. This is expected in part because the predicted log2 ratios are bounded by Equations 1 and 2 and use the averages of the three pure samples in both equations. For the subset miRNAs measured with PCR (Figure 4, panel F), the observed ratios confirm that the mixture design provides the predicted differences. Estimation of the useable region of the dynamic range based upon deviation from benchmark log2 ratios has been described for mRNA measurements using microarrays [9]. This metric relies on the subset of tissue-selective miRNAs to behave similarly to log2 ratio values derived directly from the mixture proportions. However, as shown in Figure 3, measuring the pure tissue components provides predicted log2 ratio values for every detected miRNA, and these can be used for direct comparison to the corresponding observed log2 ratios. By assessing the deviation from predicted log_2_ ratios for all observed values, the entire measurement system can be evaluated using all miRNA regardless of their level of enrichment in any single tissue component. Figure 5 shows these differences for the same data as Figures 3 and 4. As summary metrics for the overall measurement system, the median deviation value can be used as an indicator of bias and the inter-quartile range (IQR) can be used as an estimate of precision (solid and dashed horizontal lines, respectively). An IQR for each tissue-selective classification can also be determined (see Figure 6).

**Figure 4.**
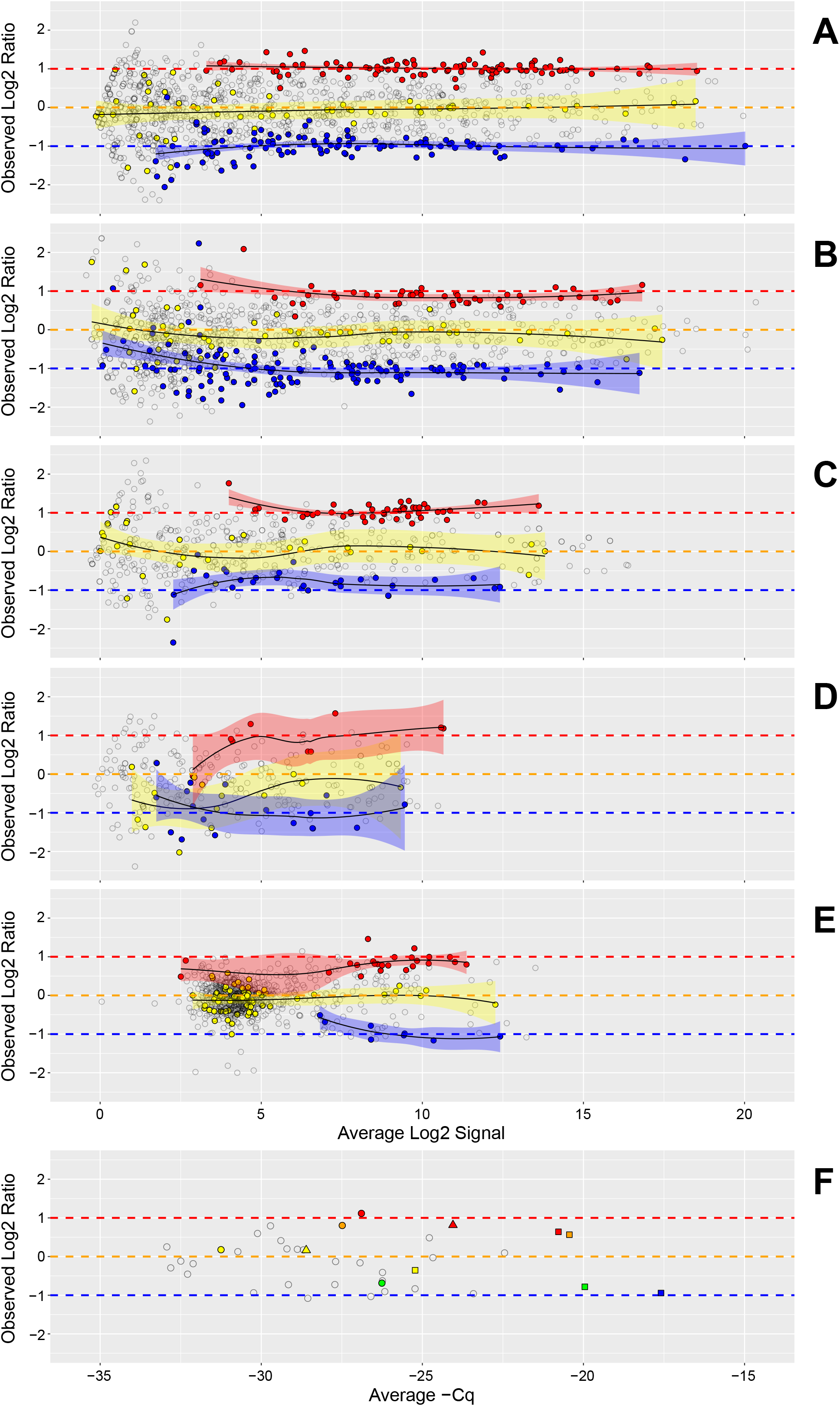
Observed distribution of log2 ratios. Panels A to F correspond to measurement processes A to F described in Results. For each datapoint in panels A to F, the difference between the Mix1 and Mix2 log2 signals (log2 ratios) is plotted against their average for each detected miRNA. Filled circles correspond observed values for miRNA that were at least 10-fold different in relative abundance compared with the other two tissues or miRNA that were approximately equal in relative abundance between placenta and brain (1-to-1): red = 10X placenta, blue = 10X brain, and yellow = 1 -to-1 (10X liver and placenta = brain). Open circles correspond to detectable, but nonselective miRNA. Red, yellow, and blue transparent bands indicate the 95 % confidence interval for the loess (locally weighted smoothing) function (black lines) for the placenta, 1 -to-1, and brain subsets, respectively. Panel F includes data from three different PCR labs: one site using multiplexed PCR (circles) and two sites using individual PCR assays (squares and triangles). Five miRNA of interest are highlighted: miR-451a (red), miR-335 (orange), miR-375 (yellow), miR-218 (green), miR-125b (blue).

**Figure 5.**
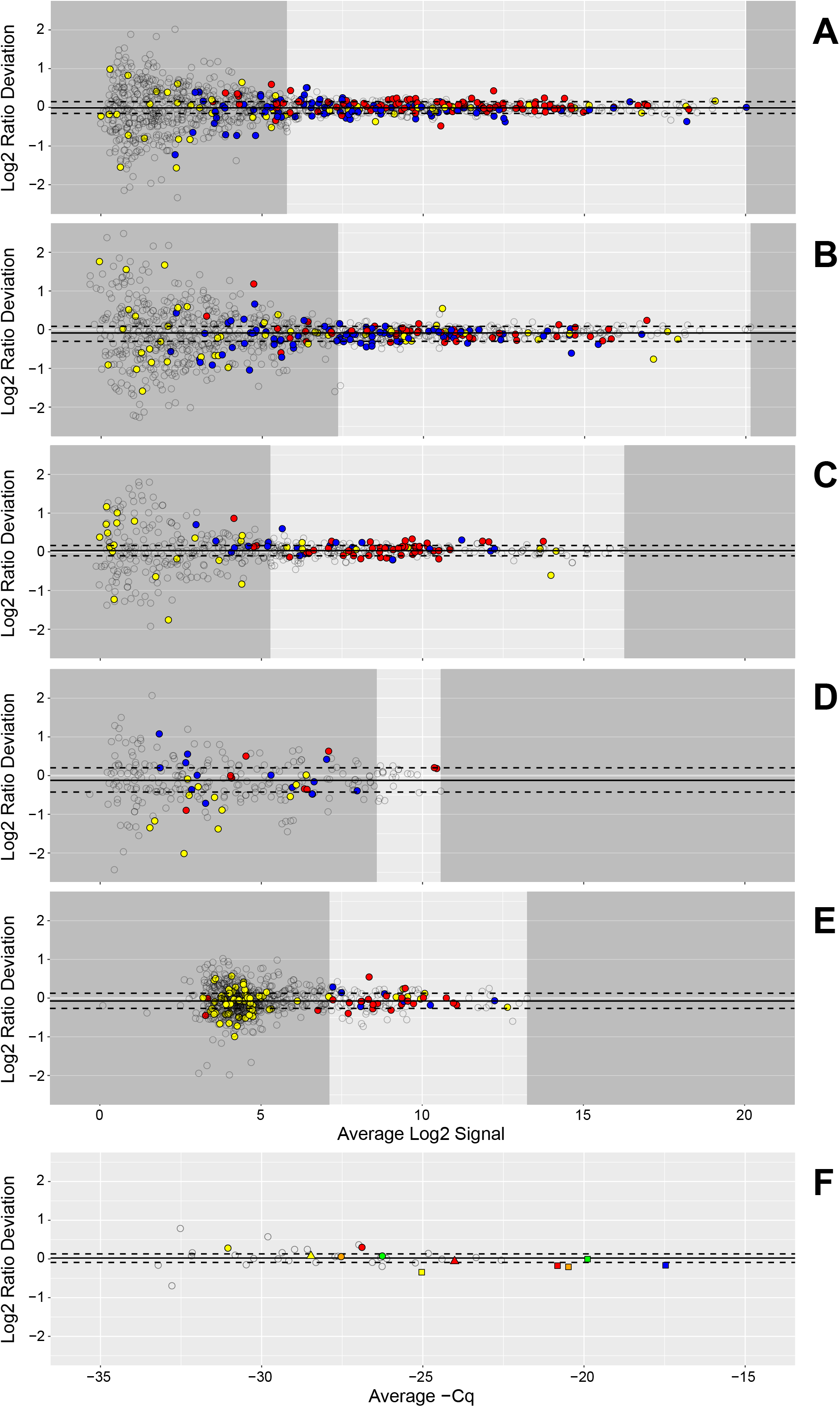
Deviation from predicted ratios as a function of dynamic range. Panels A to F correspond to measurement processes A to F described in Results. Each datapoint in panels A to F represents the difference between the observed and predicted log2 ratios plotted against the average observed and predicted log2 signal for each detected miRNA. Open circles correspond to all detectable non-selective miRNA and yellow, blue, and red filled circles correspond to 1 -to-1, brain-, and placenta-selective miRNA, respectively. The median and interquartile range (IQR) of the deviation from predicted for all detected miRNA are indicated by the solid and dashed horizontal lines, respectively. The lower limit of acceptable dispersion (determined by a user selectable deviation of ± 0.585 log2, see Results) and the maximum detectable value are indicated by the margins of the darker grey areas, respectively. Margins were not assessed in Panel F. Panel F includes data from three different PCR labs: one site using multiplexed PCR (circles) and two sites using individual PCR assays (squares and triangles). Five miRNA of interest are highlighted: miR-451a (red), miR-335 (orange), miR-375 (yellow), miR-218 (green), miR-125b (blue). *Limits and range included in the summary table of the dashboard.*

**Figure 6.**
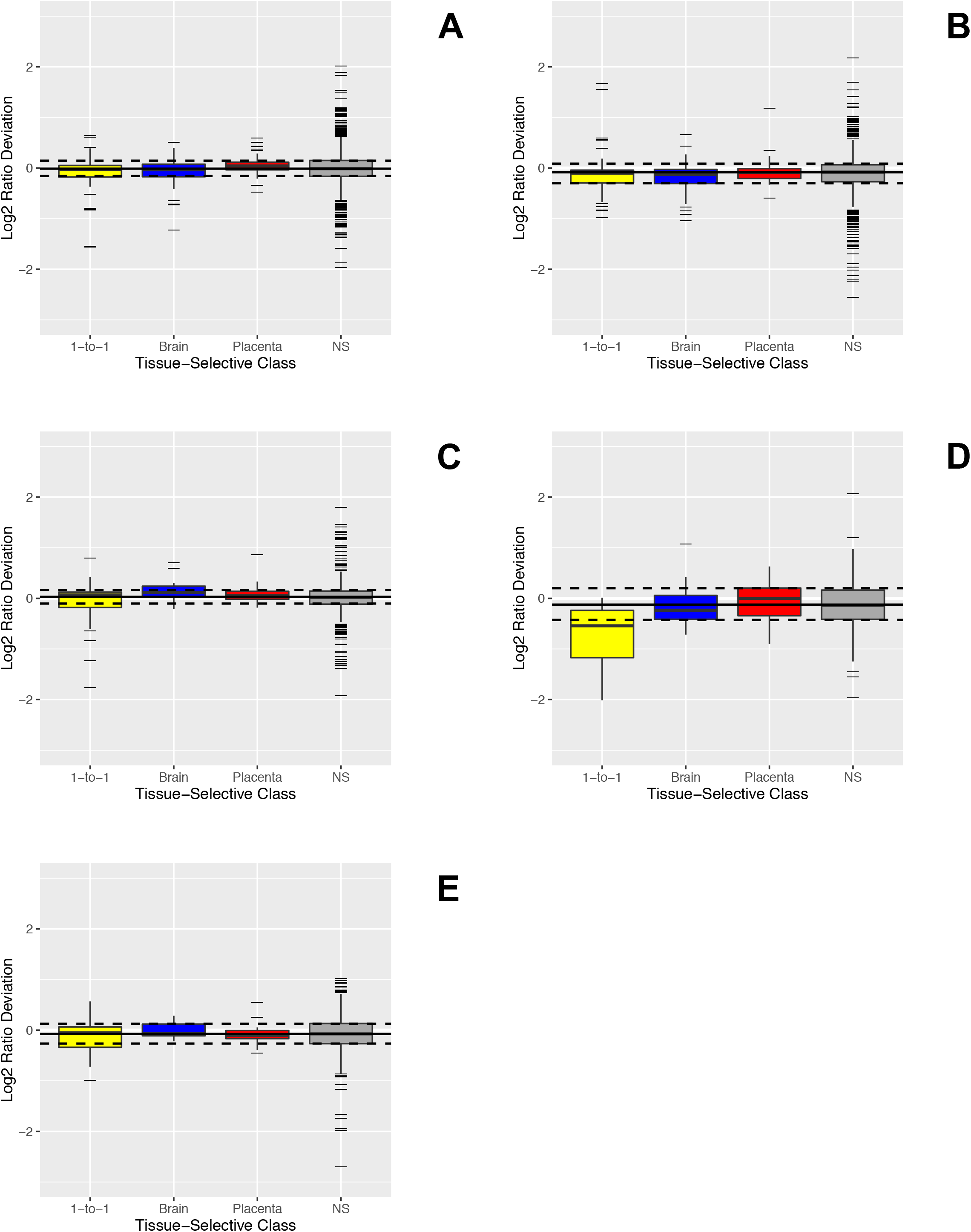
Bias and dispersion within tissue-selective classes. Panels A to E correspond to measurement processes A to E described in Results. The bias and dispersion for each tissue-selective class is shown in box (IQR) and whisker (1.5*IQR) format with outliers represented by black hash marks and median values indicated by black line (yellow = 1 -to-1, blue = brain, red = placenta, and grey = none-selective (NS)). The median and interquartile range (IQR) of the deviation from predicted for all detected miRNA are indicated by the solid and dashed horizontal lines, respectively.

It is clear that the majority of values falling outside the IQR occur at the lower end of the dynamic range. A lower limit for the useable range of the measurement system can be determined for a particular level of tolerance for deviation — for example, modeling how the distribution of deviation changes along the dynamic range and locating the lowest average log2 signal for which at least 95 % of the modeled distribution falls within one half fold-change (± 0.585 log2 difference). This lower limit and the maximum value demarcate the reliable region of the dynamic range in Figure 5. A tolerance for deviation can also be used to define the upper limit for technologies that may experience performance declines at higher signals, for example saturation in microarrays [9].

The tissue-selective subsets provide both true positive (brain and placenta) and true negative (1-to-1) classifications useful for preparing receiver-operating characteristic (ROC) curves, and the area under the curve (AUC) can be used as a summary metric [7]. For each site, the tissue-selective miRNA are ranked by p-values using a paired t-test comparison of the log2 signals for the three replicate Mix1 and Mix2 samples. Figure 7 shows the corresponding ROC curves for all detected tissue-selective data as well as the ROC curves derived from data within the reliable region of the dynamic range described in Figure 5. Within a laboratory, both the AUC and the reliable range can be used to monitor alterations in performance introduced by changes in technology, reagents, or operator experience [7–9]. However, the AUC metric is limited by the availability of true positive and true negative differences identified by the pure sample profiles, and direct comparisons between different measurement systems may not be meaningful if there is a significant difference in the number of miRNA being assessed. Range limitations and AUC values per site and round are included in Table 1 and in the metrics tables of Additional file 1.

**Figure 7.**
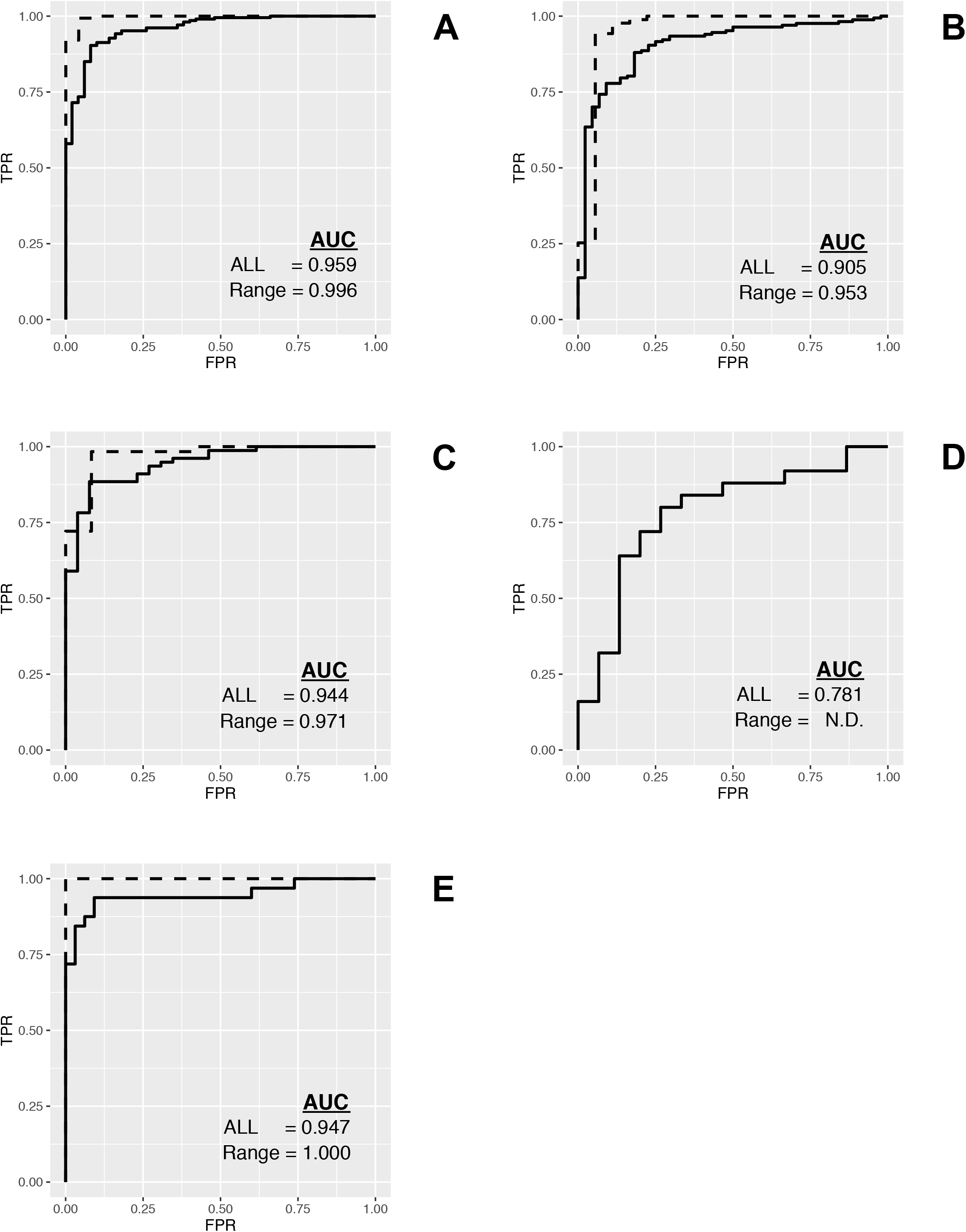
Discrimination accuracy. Panels A to E correspond to measurement processes A to E described in Results. Receiver-Operating Characteristic Curves (ROCplots), where true positives correspond to 10X placenta and 10X brain miRNA; and true negatives correspond to 1 -to-1 components using either the entire detectable range (solid line) or limited to the reliable region (dashed line). *Area under the curve (AUC) values are included in the summary table of the dashboard.*

**Table 1.**
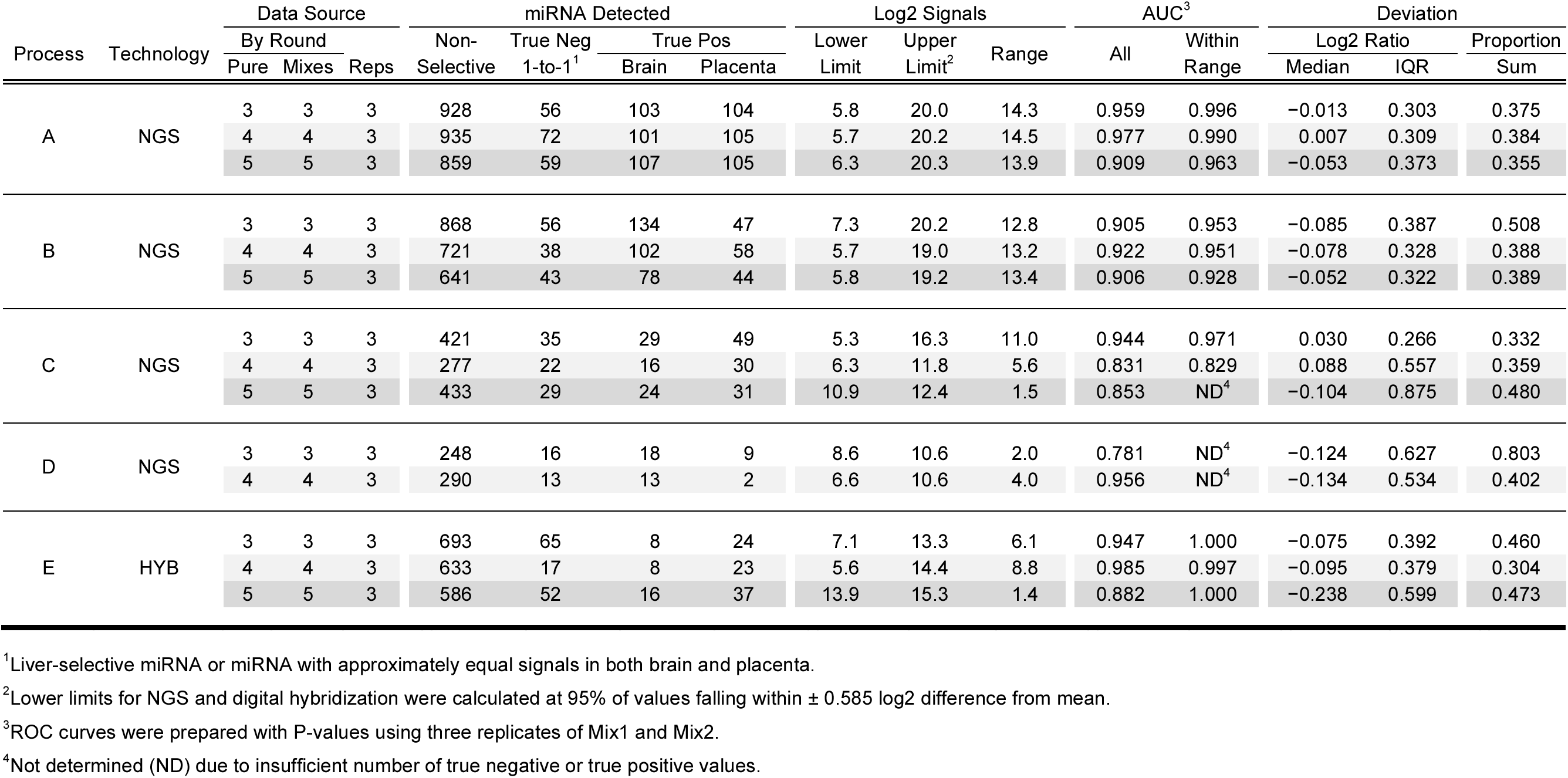
Summary metrics derived from complete reference sample sets: five samples, three replicates each, and three rounds.

### Proportion-based metrics

While ratio based metrics are useful for evaluating a sites ability to accurately detect differences between samples, evaluating the measurements for each individual mixture may provide additional information. A model can be fit based on Equations 1 and 2, and solved for the expected proportions of *Φ* (see above) given the set of signals, *S*_*i*_. Genome-scale data can confidently estimate these proportions from the collected data for each pure component and mixture. Deviations from the designed proportions can be visualized using *target* plots (Figure 8) [10]. The (x, y) coordinates for the center of each target correspond to the designed proportions and the (x, y) coordinates for the end of each line segment emanating from the center correspond to the estimated proportions of each pure tissue component in Mix1 and Mix2, respectively. The lengths of these line segments provide an indication of potential bias in the measurement process. Part of this deviation is due to intrinsic differences in the miRNA content of each pure tissue, seen in the consistency of the direction of the yellow and red lines across labs. An mRNA fraction effect has been observed for transcriptomic measurements, and a means to assess this has been developed using RNA spike-in controls simulating poly-A mRNA [18]. The proportion estimates and bias indicators for each component are included in the metrics table of the dashboard. The sum of all lengths may provide a single useful indicator and is included in the dashboard summaries and in Tables 1 to 3. The ellipses surrounding the line segment ends indicate the 95 % confidence intervals for the estimated proportions of each component in Mix1 and Mix2 and are influenced by a combination of both dispersion and detection within each tissue-selective class. Indications of poor precision are apparent in the measurement process shown in panel D in both Figure 6 and 8.

**Figure 8.**
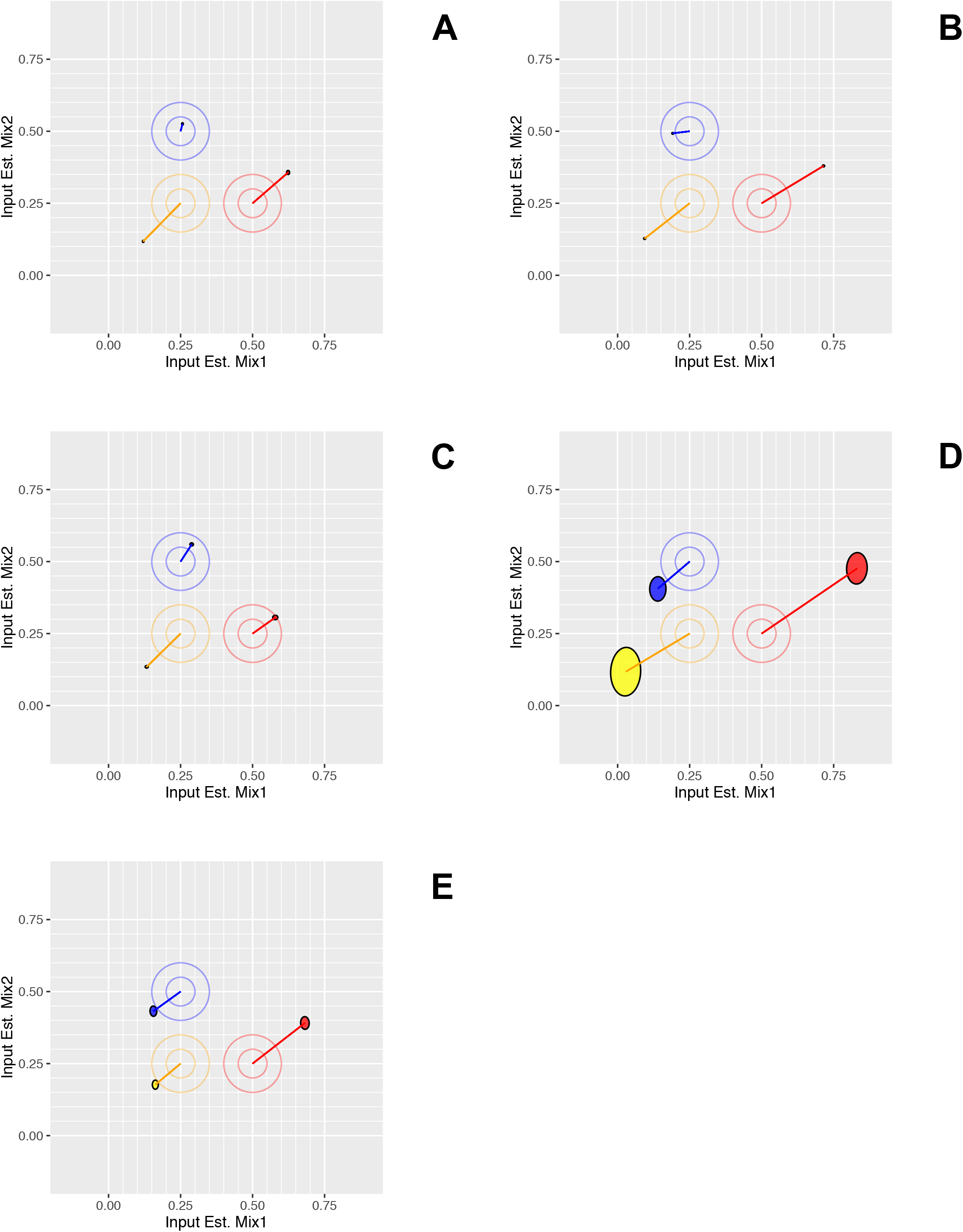
Deconvolved mixture proportions of tissue components in Mix1 and Mix2. Panels A to E correspond to measurement processes A to E described in Results. Concentric circles added to target values for emphasis. Line segments connect target values (central point of circles) to their corresponding estimates. Ellipses show the 95 % confidence interval range of the mixture proportions for the three tissue components (yellow = liver, blue = brain, and red = placenta).

### Minimizing the experimental design

It should be noted that each round of the study included three replicates each of three pure total RNAs, and three replicates each of two mixtures for a total of 15 samples, and predicted values were derived from and compared with samples processed at the same time. Figures 2 to 8 and the data in Table 1 correspond to these within round comparisons using three replicates. Modifying the experimental design to minimize the number of samples required may be accomplished it two ways.

First, pure total RNA, prepared as part of large enough batch to provide mixture samples that span several rounds may provide a sufficient baseline in the first round to allow for subsequent rounds of testing to be based on the paired mixtures alone, reducing the number of samples required for processing to six. To test this, we used the Round 3 pure tissue profiles as the baseline predicted values for comparison with the mixtures in the subsequent rounds. Metrics derived from this baseline approach are included in Table 2. Using predicted data derived from Round 3 pure samples alone neither obscures, nor distorts the differences among the measurement processes shown in Table 1. This indicates that using the paired mixtures alone, after establishing a baseline prediction, might be sufficient for monitoring processes over time. Intentional changes to a measurement process (e.g., reagent kits, instrumentation, or software) may require re-evaluation of the pure components. In this study, aliquots initially prepared as part of one large set of samples prior to Round 3 were distributed to participants every six months for Rounds 4 and 5. Reference sample stability for longer periods has not been evaluated.

**Table 2.**
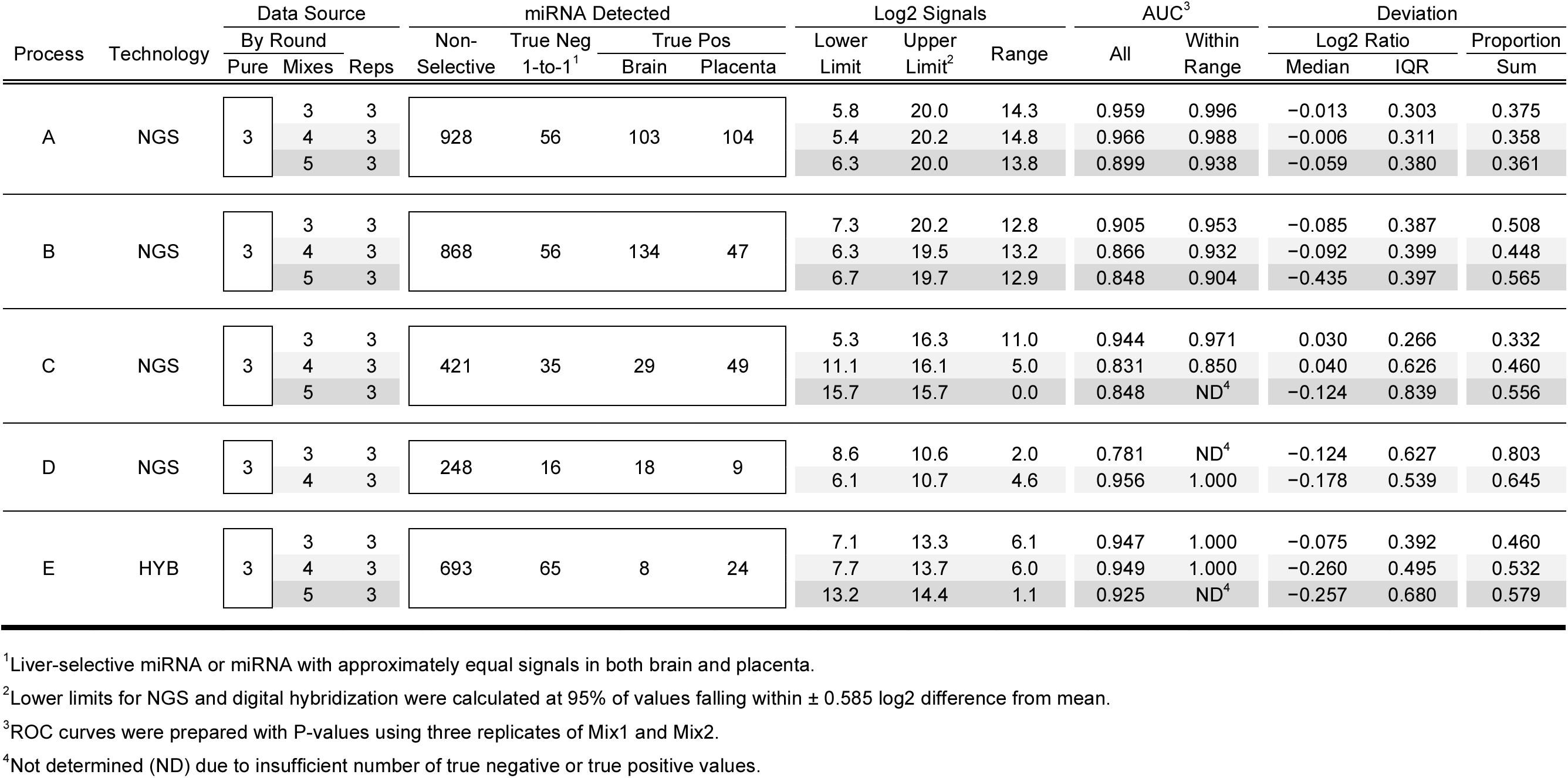
Summary metrics derived from using pure samples from Round 3 for baseline predictions.

The second approach to reducing the number of samples would be running the sample set (brain, liver, placenta, Mix1, and Mix2) without replicates. To test this we limited the analysis to the first replicate of each dataset. Metrics derived from this approach are included in Table 3. In this case, the ROC curves derived from datasets without sample replication are based on ordered ratios instead of P-values [7]. In the absence of technical replication, the resulting AUCs are lower when all tissue-selective miRNA are evaluated, the lower limit of the useable range is higher, and the IQR is increased. Therefore a consistent approach, either with or without replication, should be used when tracking a measurement process over time. It should also be noted that, in the absence of replication, a measurement failure for any one the five samples in the set would render some of the metrics indeterminable.

**Table 3.**
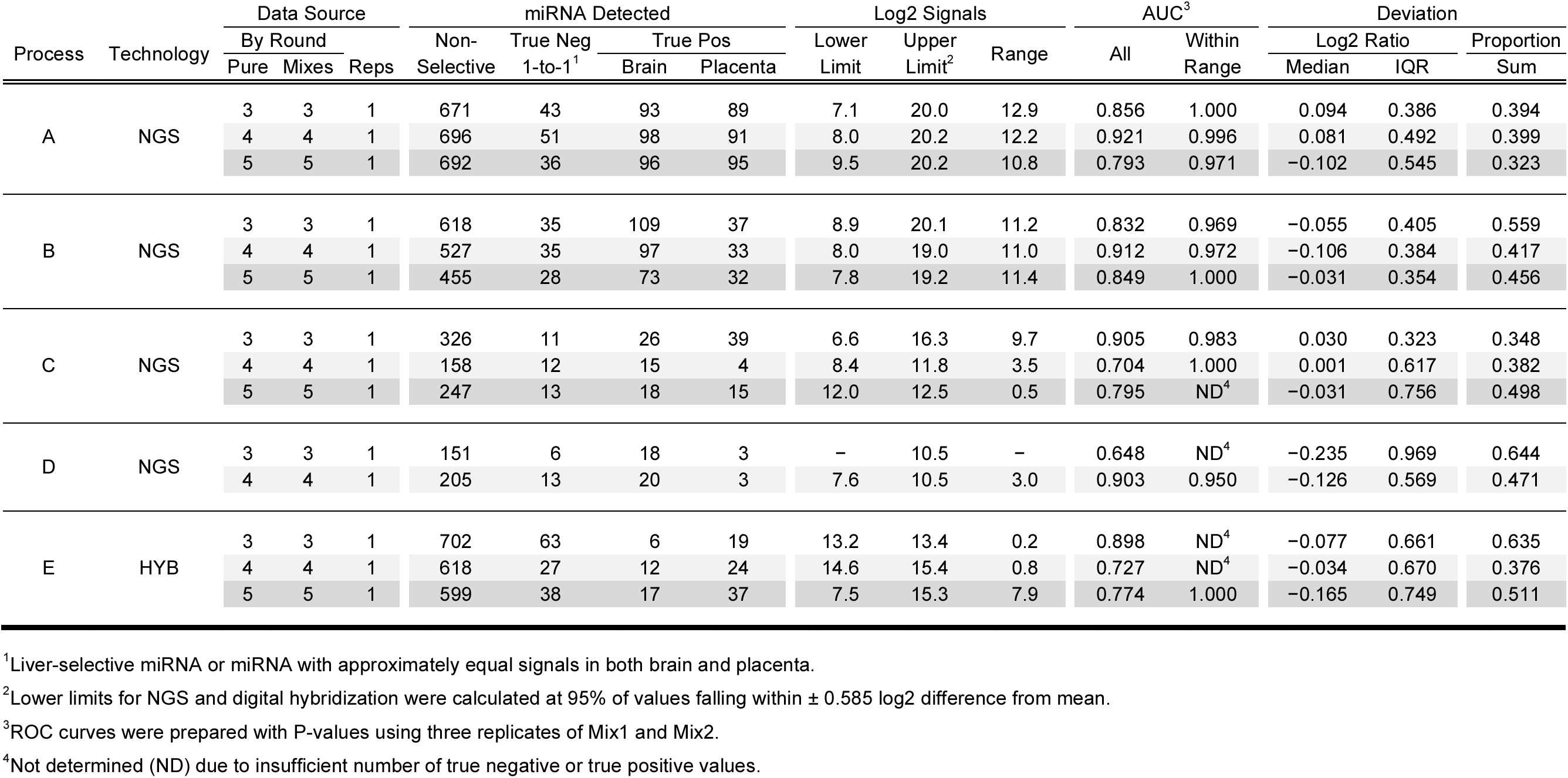
Summary metrics derived from using one replicate of each sample per round.

## Discussion

The total RNA reference sample set described here provides process controls for genome-scale measurements of miRNA that are reasonable biological mimics and provide a sufficient number of miRNAs to span the dynamic range of a measurement system. Evaluation of the deviations in ratios designed into the sample set provides a quantitative assessment of the reliable region of the measurement system. These sample sets, with or without replication and with or without baseline, can be run in parallel with or in between biomarker profiling experiments at some frequency to provide ongoing measurement assurance of the complete measurement process. An observation of poor results with the process controls (lower AUC, decreased reliable range, etc.) may indicate that profiling experiment results from a proximal timeframe may also have issues. However, achieving an acceptable result with the process controls only indicates that a *measurement process* is working well, but does not confirm experimental observations on the samples under study.

This study was designed to evaluate the reference samples and develop associated metrics, and is not intended as a platform comparison. The results presented are from those sites that accepted samples for the rounds that included mixtures and subsequently provided data for analysis. Additional details for each measurement process are available in Additional file 2 as outlines of the protocols in place at the labs; they are not intended to be reproducible resources. This study demonstrates of the utility of these mixture samples and associated metrics to evaluate technical performance of any genome-scale measurement process, the methods and protocols are incidental to the study presented here. A and B are the only measurement processes using the same platform. Measurement processes C and D were performed at a single site using two different platforms (both different from the platform used in A and B). Measurement processes E and F are unique sites and platforms.

The current dataset collection provides a range of performance and demonstrates that the samples, the visualizations presented in Figures 2 to 8, and the summary metrics shown in Table 1 can be used to discern differences in performance. Systematic application of the samples, metrics, and methods described here can enable evaluation and optimization of both laboratory and measurement platform performance. Evaluating the relationship between protocols used at different sites and observed performance may also be useful to identify key parameters for optimization.

To promote periodic self-assessment of genome-scale measurement system performance, a web-based version of the analysis pipeline has been implemented as part of the EDRN Informatics Center [https://xxxx.xxx.xxx.xxx/xxxxxxxxx]. Dashboard views of the results can also be generated online (see Additional file 3 for instructions). Visitors to the site may view descriptions of available reference samples or download a protocol on how to prepare them in their own laboratory [14] (a brief description is available in Methods). Visitors may also view or download publicly available datasets and results. Current participants can add to their datasets and compare the new results to prior datasets to assess individual site performance over time. For new sites interested in assessing their genome-scale profiling workflows, information about registration and availability of EDRN prepared reference sample sets is provided at the EDRN website.

## Conclusions

Metrics and visualizations derived from mixture samples are well suited for assessing performance of genome-scale measurement systems used to identify differentially regulated miRNAs. They are made from biological materials similar to those studied by biomarker profiling laboratories and provide a sufficient number of differentially expressed miRNAs with predictable ratios to serve as benchmarks. Implementing these metrics and visualizations as part of an online resource offers laboratories the opportunity to evaluate and optimize their discovery process.

## Additional files

Additional file 1: Dashboard views of measurement processes A — E from Rounds 3-5, using three replicates (as PDF).

Additional file 2: Protocols for measurement processes A — F (as PDF).

Additional file 3: Instructions for using measurement assurance pipeline online (as PDF).

## Declarations

## Abbreviations

NIST: National Institute of Standards and Technology
NCI: National Cancer Institute
NASA: National Aeronautics and Space Administration
JPL: Jet Propulsion Laboratory
EDRN: Early Detection Research Network
Caltech: California Institute of Technology
RT-PCR: reverse transcription polymerase chain reaction
NGS: next-generation sequencing
HYB: hybridization

## Ethics approval and consent to participate

Not applicable.

## Consent for publication

Not applicable.

## Availability of data and material

The datasets supporting the conclusions of this article are included within the article, its Additional files, and online at [https://xxxx.xxx.xxx.xxx/xxxxxxxxx].

**Funding**

This research was supported by an NCI-EDRN joint NIST-Biochemical Science Division Interagency Agreement and NCI-EDRN Grant numbers: U01CA214182, U01CA214195, U24CA115091. Part of the work was performed at JPL/Caltech under the contract to NASA, and at the Center for Data-Driven Discovery, Caltech.

## Authors’ contributions

PSP, LS, and MLS designed the study. PSP, LKV, and MLS developed the reference samples. LKV, AS, GL, ACG, HIP, CG, SMD, KK, SAS, DK, KVKJ, ACL, KLT, and BAR acquired and processed the data. PSP, SPL, and JRP developed metrics and visualizations. PSP, AAM, LC, SCK, HK, and DJC developed the website. PSP drafted the manuscript. SPL, JRP, AAM, and LC contributed manuscript sections. All authors participated in the revision process and provided final approval.

## Acknowledgements

Not applicable

## Disclaimer

Certain commercial entities, equipment or materials may be identified in this document in order to describe an experimental procedure or concept adequately. Such identification is not intended to imply recommendation or endorsement by the National Institute of Standards and Technology, nor is it intended to imply that the entities, materials or equipment are necessarily the best available for the purpose.

## Methods

### Mixture design

Human Brain Reference RNA (Cat. No. AM6050), Human Liver Total RNA (Cat. No. AM7960), and Human Placenta Total RNA (Cat. No. AM7950) was obtained from Ambion (Thermo Fisher Scientific). Manufacturers stock solutions of 1 μg/μl were verified on a Qubit (Thermo Fisher Scientific). If necessary, stock solutions of pure tissue components (same lot numbers) were combined to provide a sufficient volume of identical material prior to distribution. Prior to mixing, an adequate portion of stock solutions are set aside for pure tissue aliquots. The remaining liver, brain, and placenta stocks were then mixed by volume using the proportions of 1:1:2 and 1:2:1 for Mix1 and Mix2, respectively. These five samples (three neat tissues and two mixtures) were then divided into aliquots. Three replicates of each sample were distributed to participants as a numbered blinded set of 15 tubes. A general method for the preparation of two mixtures of total RNA (Mix1 and Mix2) derived from three different pure total RNA sources (RNA1, RNA2, and RNA3) from either commercially available or laboratory prepared total RNA is also available [14]. This protocol allows labs to recreate previously measured sample designs for comparison or to generate new sample designs with different components and/or mixture formulations.

### Sample handling and analysis

Each laboratory used its routine protocol for miRNA biomarker detection and evaluations. Individual protocols are available online as part of the data repository, and included in Additional file 2.

